# Rad51 inhibition sensitizes non-replicating quiescent cells to UVB radiation and transcription stress

**DOI:** 10.1101/2022.09.20.508657

**Authors:** Saman Khan, M. Alexandra Carpenter, Michael G. Kemp

## Abstract

DNA damage induced by environmental, occupational, and chemotherapeutic compounds lead to a variety of cellular responses that are potentially impacted by the proliferation status of the cell. Using small molecule inhibitors of various DNA doublestrand break (DSB) repair pathways in non-replicating, quiescent human cells exposed to UVB radiation, we unexpectedly observed a major role for the recombination protein Rad51 in promoting cell survival. In contrast to a previous report indicating a requirement for nucleotide excision repair (NER) in DSB formation after UV exposure in quiescent cells, we observed DSB formation and Rad51 function to be independent of NER. Moreover, our analyses of DNA damage response kinase signaling in quiescent cells identified protein substrates that were either dependent or independent of both NER and apoptotic signaling. Finally, we observed that Rad51 inhibition sensitized nonreplicating quiescent cells to diverse genotoxic stressors, including those inducing DNA-RNA hybrids and inhibiting transcription. Thus, these findings clarify the mechanisms by which DSBs arise in non-replicating cells and highlight the important role of Rad51 in promoting quiescent cell survival in response to general genotoxic stress.

**Summary statement:** DNA double strand breaks are generated independent of nucleotide excision repair in UV-irradiated quiescent cells and require the recombination protein Rad51 to promote cell survival.

## Introduction

Much of our understanding of cellular DNA damage response pathways has been derived from model systems (yeast, Xenopus egg extracts, cancer cell lines) in which DNA replication and cell cycle progression are major contributors to both the mechanism by which the damage is sensed and the downstream effects of the damage on the cell. A classic example involves exposure to UV radiation. In systems in which chromosomal DNA synthesis is operational, UV photoproducts and other bulky DNA adducts are thought to uncouple the coordination between DNA helicases and DNA polymerases to generate single-stranded DNA (ssDNA) and primer-template junctions that lead to the recruitment and activation of the ATR protein kinase (Byun et al., 2005; MacDougall et al., 2007), which regulates cell cycle phase transitions, replication fork stability, and replication origin firing (Perry and Kleckner, 2003; Saldivar et al., 2017). Early studies of ATR kinase signaling therefore concluded that ATR activation in response to UV radiation required replication stress (Lupardus et al., 2002; Ward et al., 2004).

However, contact inhibition and serum starvation to generate G0/quiescent cells and flow cytometry to specifically examine G1 phase cells were subsequently used by several groups to show that UV radiation could induce the phosphorylation of ATR substrate proteins such as the histone variant H2AX in non-S phase cells in a manner dependent on ATR and nucleotide excision repair (NER) but independent of DSB formation (Hanasoge and Ljungman, 2007; Marini et al., 2006; Marti et al., 2006; Matsumoto et al., 2007; O’Driscoll et al., 2003). One report also indicated a role for the base excision repair protein Ape1 in ATR-mediated H2AX phosphorylation in UV-irradiated quiescent cells (Vrouwe et al., 2011). In contrast, other groups have reported robust phosphorylation of both the tumor suppressor protein p53 (Derheimer et al., 2007; Ljungman et al., 2001) and the ATR kinase (Kemp, 2017) in cells deficient in NER that may be due to RNA polymerase II stalling and transcription stress (Derheimer et al., 2007; Ljungman et al., 2001). Thus, there are likely more than one mechanism by which ATR can become activated in non-S phase cells.

However, the most widely understood and accepted model for ATR kinase signaling in response to UV radiation in G0/G1 cells is that the ssDNA gaps generated by the NER machinery are then enlarged by the nuclease Exo1 to result in a DNA substrate of sufficient length for recruitment and activation of ATR (Lindsey-Boltz et al., 2014; Sertic et al., 2011). Furthermore, additional work demonstrated that in the absence of efficient gap filling synthesis (Matsumoto et al., 2007), including by translesion synthesis DNA polymerases (Sertic et al., 2018), DNA double-strand breaks could arise (Sertic et al., 2018; Wakasugi et al., 2014) and lead to the activation of the ATM kinase (Wakasugi et al., 2014). Thus, the current model for UV-dependent DSB formation and ATM activation in non-replicating quiescent cells is that it requires NER.

In this study, we re-examined the requirement for NER in the generation of DSBs in UV-irradiated quiescent cells by using CRISPR/Cas9 genome editing to knockout the core NER protein XPA in human keratinocytes to allow for analyses of DNA damage responses in NER-proficient and -deficient cells with an identical genetic background. Though these XPA-knockout cells are fully deficient in UV photoproduct removal and DNA repair synthesis, we observed that DSBs readily formed in these cells following UVB exposure. Moreover, through a small pharmacological screen of DSB repair inhibitors, we unexpectedly found that the homologous recombination protein Rad51 promotes the survival of both wild-type and XPA knockout quiescent cells after exposure to UV and a variety of different genotoxic agents, including compounds that induce transcription stress. Together, these results support a non-NER-dependent pathway by which DSBs can arise in quiescent UV-irradiated cells and implicate a critical function for Rad51 in mediating the survival of non-replicating quiescent cells exposed to DNA damaging agents.

## Results

### Characterization of a model cell culture system of quiescence

A common approach to generate a population of non-replicating, quiescent human cells is to grow cells to confluence and then incubate for a few days in culture medium with a low concentration of serum. To better characterize this commonly used approach (Hanasoge and Ljungman, 2007; Kemp, 2017; Kemp and Sancar, 2016; Marini et al., 2006; Matsumoto et al., 2007; O’Driscoll et al., 2003; Sertic et al., 2011; Sertic et al., 2018; Wakasugi et al., 2014), we performed flow cytometric analysis of propidium iodide-stained HaCaT keratinocytes growing in a sub-confluent proliferating state with medium containing 10% serum or in cells grown to confluence for 2-3 days and then incubated for 2-3 additional days in medium containing 0.5% serum to reach a quiescent state. As shown in **Fig. 1A**, whereas the proliferating cells displayed a significant population of cells with an intermediate S phase DNA content between the characteristic G1 and G2 peaks, few cells were observed in this location when in the quiescent state. Furthermore, direct detection of ongoing DNA synthesis by EdU labeling showed that less than 1% of cells are synthesizing significant amounts of DNA (**Fig. 1B**). However, because cells with a G2 DNA content are present in this population of cells (Fig. 1A), there are likely a small number of cells that move into and through S phase and accumulate at G2/M under these culture conditions. MTT assays showed that treatment with the DNA replication inhibitors gemcitabine or hydroxyurea caused reduced cell number/proliferation in cells in the proliferating state but not in cells the quiescent state **(Fig. 1C, D**). We conclude that very few HaCaT cells are actively synthesizing DNA when grown to confluence and maintained in low serum, which provides further validation for this cellular model of quiescence.

**Fig. 1.**
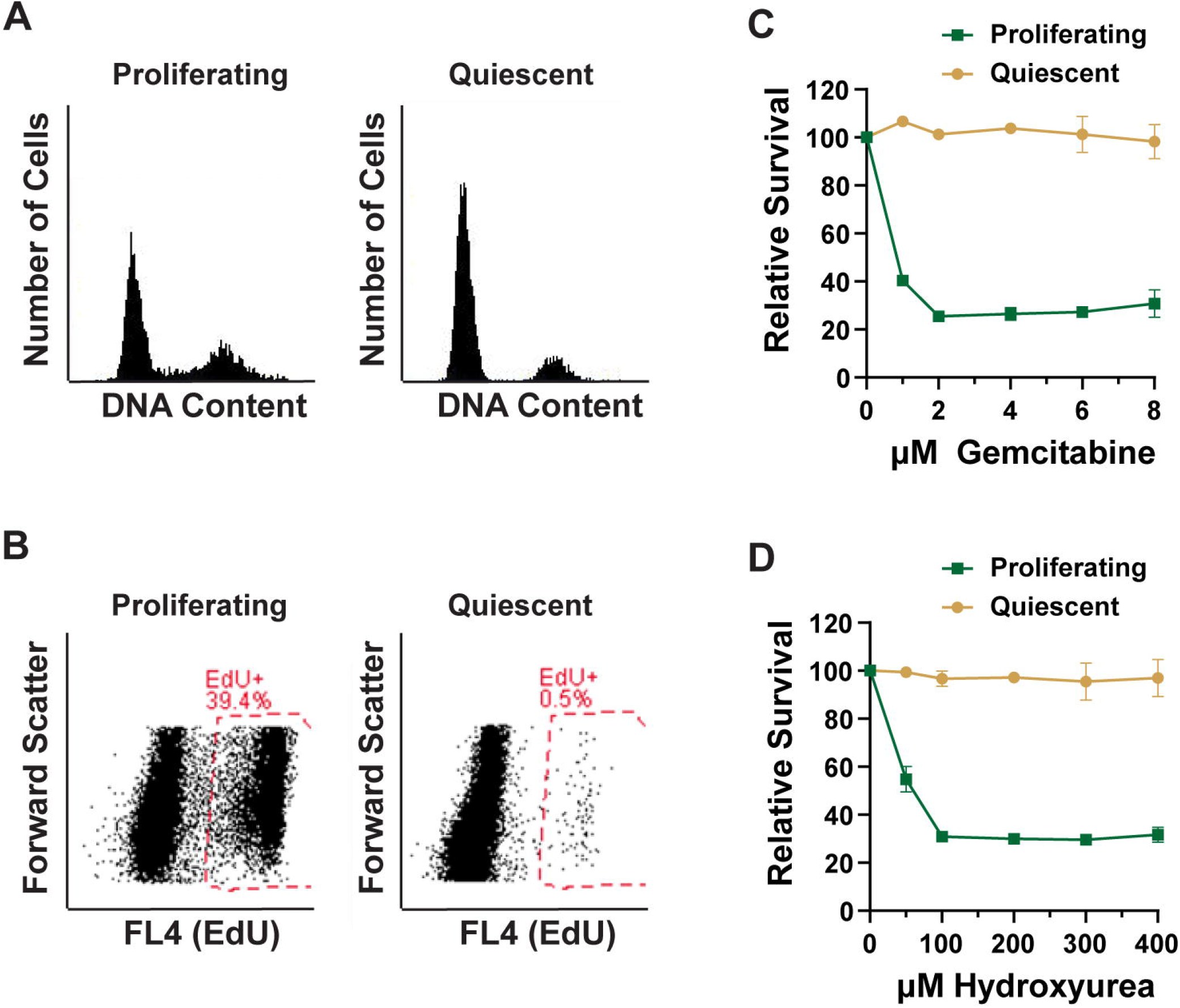
Characterization of quiescent cells. **(A)** Flow cytometric analysis of propidium iodide-stained HaCaT cells maintained in a sub-confluent state in 10% serum (Proliferating) or confluent state for 2-3 days in 0.5% serum (Quiescent). **(B)** Proliferating and quiescent HaCaT cells were pulsed with 10 μM EdU for 2 hr before analysis by flow cytometry and quantitation of the percentage of EdU-positive cells. **(C, D)** Proliferating and quiescent HaCaT cells were treated with the indicated concentrations of gemcitabine or hydroxyurea for 3 days. MTT assays were performed to examine relative cell number/survival.

### Rad51 promotes UVB survival in quiescent cells

A previous study using telomerase-immortalized fibroblasts from a xeroderma pigmentosum (XP) patient and telomerase-immortalized testicular teratoma cells from a normal, NER-proficient individual maintained in a quiescent state concluded that DSB generation and ATM kinase signaling after UV exposure was dependent on NER (Wakasugi et al., 2014). How these DSBs are repaired in these cells was not examined, but the DNA damage response protein kinase ATM was shown to be important for cellular recovery. A schematic of NER-dependent DSB formation and possible modes of repair is provided in **Fig. 2A** along with several small molecule inhibitors that have identified that target different DSB repair proteins. Quiescent HaCaT cells were treated with the indicated pharmacological agent before exposure to 0 or 400 J/m^2^ of UVB radiation and measurement of relative cell survival 3 days later using an MTT assay **(Fig. 2B**). Treatment of cells with spironolactone (SP), which induces the proteolytic degradation of the essential NER protein XPB (Alekseev et al., 2014; Kemp et al., 2019) resulted in little cell survival, thus demonstrating the well-known importance of NER in the cellular response to UVB radiation. Consistent with previous studies (Shaj et al., 2020; Wakasugi et al., 2014), inhibition of the ATM, ATR, and DNA-PK protein kinases partially sensitized the cells to UVB radiation.

**Fig. 2.**
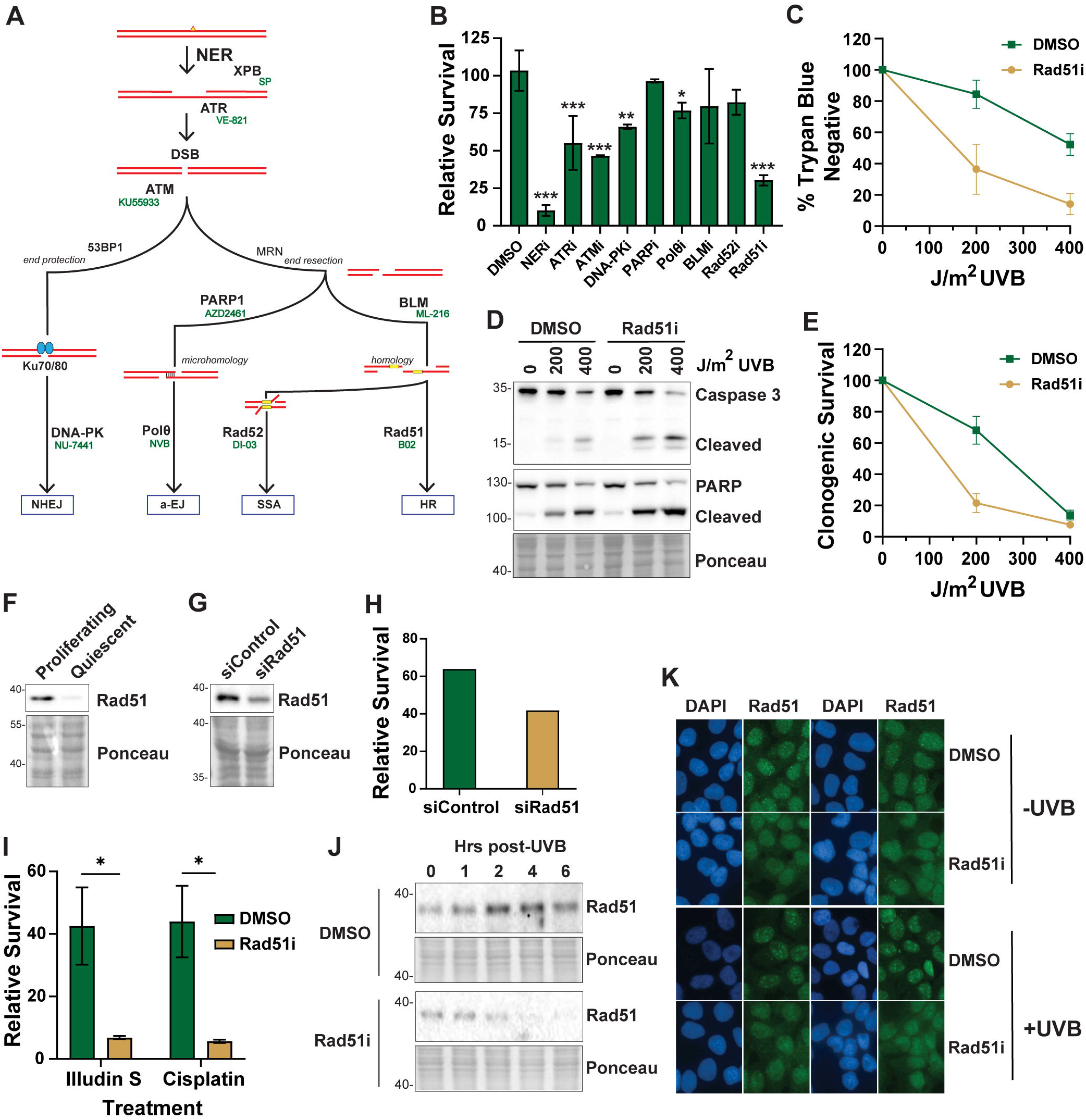
Rad51 promotes UVB survival in quiescent cells. **(A)** Schematic of DSB repair pathways activated in response to NER-dependent DSB formation. Small molecule inhibitors targeting the indicated DNA repair protein are indicated. **(B)** Relative cell survival as measured by MTT assay 3 days after treatment of quiescent HaCaT cells with the indicated inhibitor and exposure to 400 J/m^2^ UVB radiation. One-way ANOVAs were used to compare the relative survival of the different treatment groups from at least two independent experiments (*, p<0.05; **, p<0.01; ***, p<0.001). **(C)** Quiescent HaCaT cells were treated with vehicle (0.1% DMSO) or 10 μM Rad51 inhibitor (Rad51i B02) for 30 min before exposure to the indicated fluence of UVB radiation. Cell survival was measured by trypan blue staining 24 hr later, and the percentage of trypan blue-negative cells are indicated. **(D)** Cells were treated as in (C) except that whole cell lysates were prepared 12 hr after UVB exposure for immunoblot analysis of apoptotic signaling. **(E)** Clonogenic survival of quiescent HaCaT cells treated with vehicle or Rad51i and exposed to UVB. Cells were re-plated in normal medium 4 days later and then stained and counted 10-14 days later. **(F)** Immunoblot analysis of Rad51 protein levels in proliferating and quiescent HaCaT cells. **(G)** Immunoblot analysis following transfection with Control (siControl) siRNA or a pool of siRNAs targeting Rad51 (siRad51). **(H)** Relative cell survival of quiescent cells following knockdown of Rad51 and exposure to UVB. **(I)** Relative survival was determined by MTT assay in quiescent HaCaT cells treated with DMSO or Rad51i and then exposed to 400 nM Illudin S or 60 μM cisplatin for 3 days. **(J)** Quiescent HaCaT cells treated with DMSO or the Rad51i were exposed to 200 J/m^2^ UVB radiation and then fractionated to enrich for chromatin-associated proteins, which was examine by immunoblotting. **(K)** Immunofluorescence microscopy was used to examine Rad51 nuclear staining in quiescent HaCaT cells in the absence and presence of UVB exposure and after treatment with the Rad51i B02.

Somewhat unexpectedly, we observed a striking sensitizing effect in cells treated with the Rad51 recombinase inhibitor (Rad51i) B02. This latter compound has been shown to prevent Rad51 association with ssDNA (Huang et al., 2012) and therefore inhibit Rad51 function in homologous recombination (HR). Although Rad51 function and HR are predominantly thought to occur during the S and G2 phases of the cell cycle when homologous templates have been generated during S phase, we were able to detect low levels of Rad51 protein in quiescent cells (**Fig. 2F**). Furthermore, several recent reports have demonstrated functions for Rad51 in G0/G1 phase in either the repair of artificially generated DNA damage at specific genomic regions site-specific DSBs (Crefcoeur et al., 2017; Wei et al., 2015; Yilmaz et al., 2021) or under conditions that allow for DNA end resection (Chen et al., 2021; Orthwein et al., 2015; Xie et al., 2020).

These sensitizing effects of Rad51 inhibition in UVB-irradiated quiescent cells was confirmed by measurements of cell survival by trypan blue exclusion 24 hr post-UVB exposure (**Fig. 2C**) and by examination of apoptotic signaling (**Fig. 2D**). Moreover, MTT assays demonstrated that the Rad51i B02 also sensitized quiescent U2OS osteosarcoma cells to the acute effects of UVB radiation (**Fig. S1**). Furthermore, longterm cell viability/recovery assays that involved re-plating the treated quiescent cells in normal growth medium several days after UVB exposure and inhibitor withdrawal to allow for cell division further showed that Rad51 inhibition in the quiescent state negatively impacted long-term clonogenic survival (**Fig. 2E**). Lastly, to validate the pharmacological targeting of Rad51 with a genetic approach, we transfected HaCaT cells with an siRNA pool targeting Rad51, lowered the serum concentration in the media to induce a quiescent state, and then performed MTT assays 3 days after UVB exposure. Rad51 protein expression was partially reduced by the Rad51 siRNA transfection (**Fig. 2G**) and was associated with less survival after UVB exposure (**Fig. 2H**).

We next wished to determine if Rad51 plays a role in promoting quiescent cell survival in response to other UV mimetic DNA damaging agents. As shown in **Fig. 2I**, treatment of quiescent cells with either Illudin S, which induces bulky DNA adducts that solely are repaired by transcription-coupled NER (Jaspers et al., 2002), or with the platinating anti-cancer drug cisplatin, were similarly sensitized by treatment with the Rad51 inhibitor.

To further confirm a biochemical function for the Rad51 protein in quiescent cells, we fractionated cells following UVB exposure to enrich for chromatin-associated proteins. As shown in **Fig. 2J**, Rad51 levels on chromatin increased 1-4 hr after UVB exposure and then began to decline. Rad51 localization was further analyzed by immunofluorescence microscopy in non-irradiated and UVB-irradiated quiescent HaCaT cells. As shown in **Fig. 2K**, small Rad51 nuclear foci could be observed even in the absence of UVB exposure. However, exposure of the cells to UVB radiation resulted in the formation of moderately larger nuclear regions of Rad51 staining (**Fig. 2K**). Importantly, both accumulation on chromatin and foci formation were sensitive to the Rad51 inhibitor, which blocks Rad51 binding to DNA (Huang et al., 2012).

### Rad51 functions independently of NER to promote cell survival in UVB-irradiated quiescent cells

Because a previous study reported that DSBs arise in UV-irradiated cells in an NER-dependent manner (Wakasugi et al., 2014), we next wanted to determine if Rad51’s pro-survival function was impacted by the loss of NER. We therefore used CRISPR/Cas9 genome editing to disrupt exon 2 of the XPA gene in human HaCaT keratinocytes and then characterized the resulting cells. Mutation of the XPA locus was validated by DNA sequencing (**Fig. S2**). Loss of XPA expression was shown by RT- qPCR (data not shown), western blotting (**Fig. 3A**), and by acute UVB survival assays (**Fig. 3B**). Stable expression of a FLAG-tagged XPA construct in these XPA-KO cells fully restored UVB sensitivity to that found in WT HaCaT cells (**Fig. 3B**), which shows that the UVB sensitivity in XPA-KO cells was solely due the loss of XPA. As shown in **Fig. 3C**, quiescent XPA-KO cells were also more sensitive to UVB radiation than their WT counterparts as measured by MTT assay 3 days after UVB exposure.

**Fig. 3.**
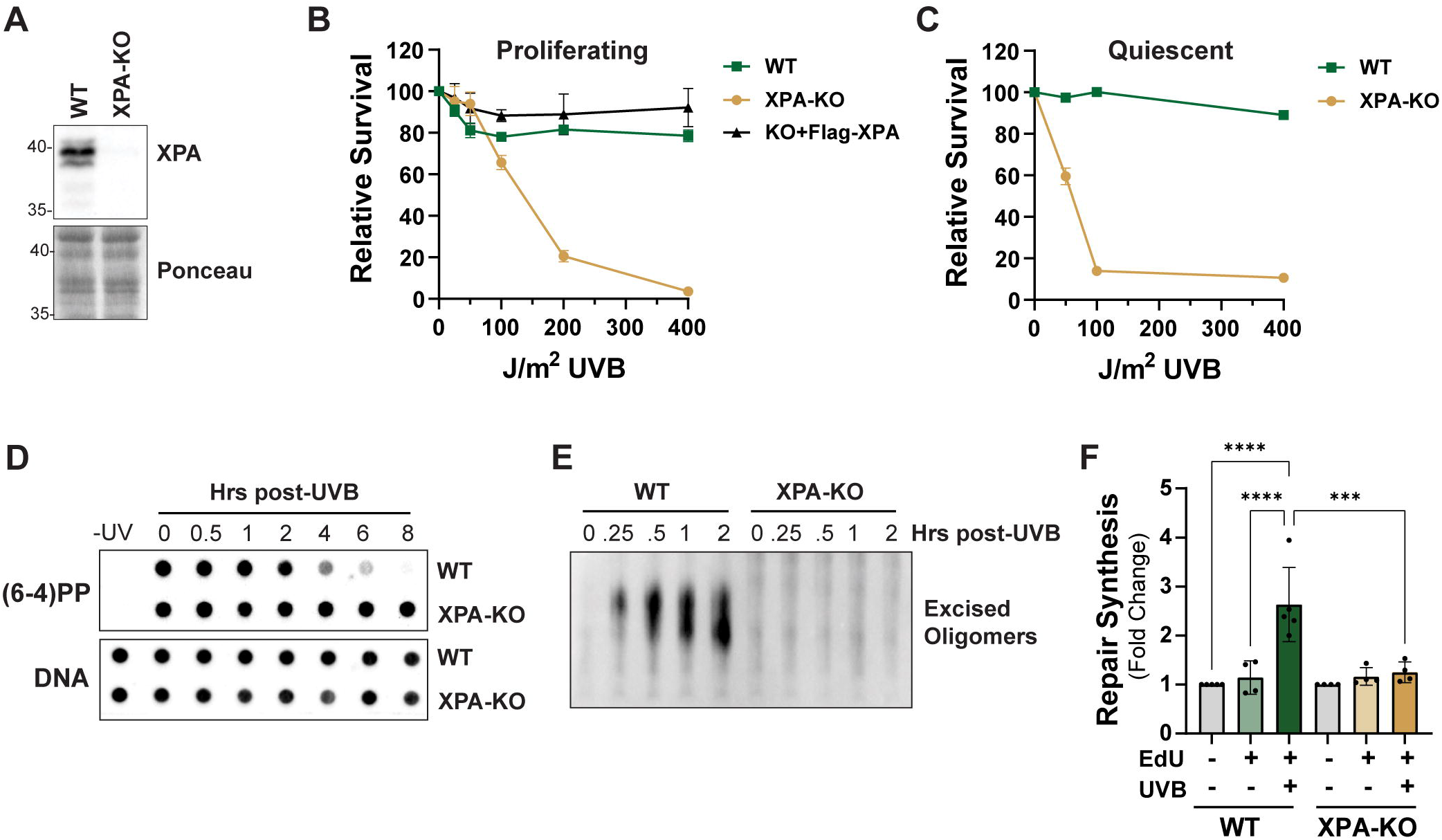
Characterization of XPA-knockout HaCaT cells. **(A).** Immunoblot analysis of XPA expression in wild-type (WT) and XPA-knockout (XPA-KO) HaCaT cells. **(B)** MTT assays were used to examine cell survival 3 days after exposure of sub-confluent cells to the indicated fluences of UVB radiation. KO+Flag-XPA indicates an XPA-KO cell line stably expressing Flag-XPA. **(C)** Relative UVB survival of non-replicating, quiescent WT and XPA-KO HaCaT cells. **(D)** DNA immunoblot analysis of (6-4)PP genomic DNA content at the indicated time points after exposure to 200 J/m^2^ UVB radiation. €**(E)** Examination of the excised DNA oligonucleotide products of NER at the indicated time points after exposure to 200 J/m^2^ UVB. **(F)** Quiescent WT and XPA-KO HaCaT cells were exposed to 200 J/m^2^ UVB and then pulsed with 10 μM EdU for 2 hr to monitor DNA repair synthesis by flow cytometry. The relative fluorescence of non-S phase cells was calculated from 4 independent experiments (one-way ANOVA; ***, p<0.001; ****, p<0.0001).

Consistent with the role of XPA in NER, XPA-KO cells were unable to remove UVB photoproducts as measured by immunodot blot analysis of (6-4)PPs in genomic DNA (**Fig. 3D**) and an assay that directly detects the excised oligonucleotide products of NER (Hu et al., 2013; Song et al., 2017) (**Fig. 3E**). Finally, flow cytometric examination of examine EdU labeling in non-S phase cells similarly showed that the quiescent XPA-KO cells were fully deficient in DNA repair synthesis (**Fig. 3F, Fig. S3**).

Having validated these XPA-KO HaCaT cells as deficient in NER, we expected to find that DSBs would not be generated following UVB exposure as was previously reported (Wakasugi et al., 2014). However, analysis of DSB formation by neutral Comet assay 2 hr after UVB exposure showed similar results in both the WT and XPA-KO quiescent HaCaT cells (**Fig. 4A**). A summary of the quantitation of the Comet tail moments is shown in **Fig. 4B**, which revealed that the only significant differences that were observed were between the non-irradiated and UVB-irradiated samples. No significant differences were found between the WT and XPA-KO cells in either the non- or UVB-irradiated cells.

**Fig. 4.**
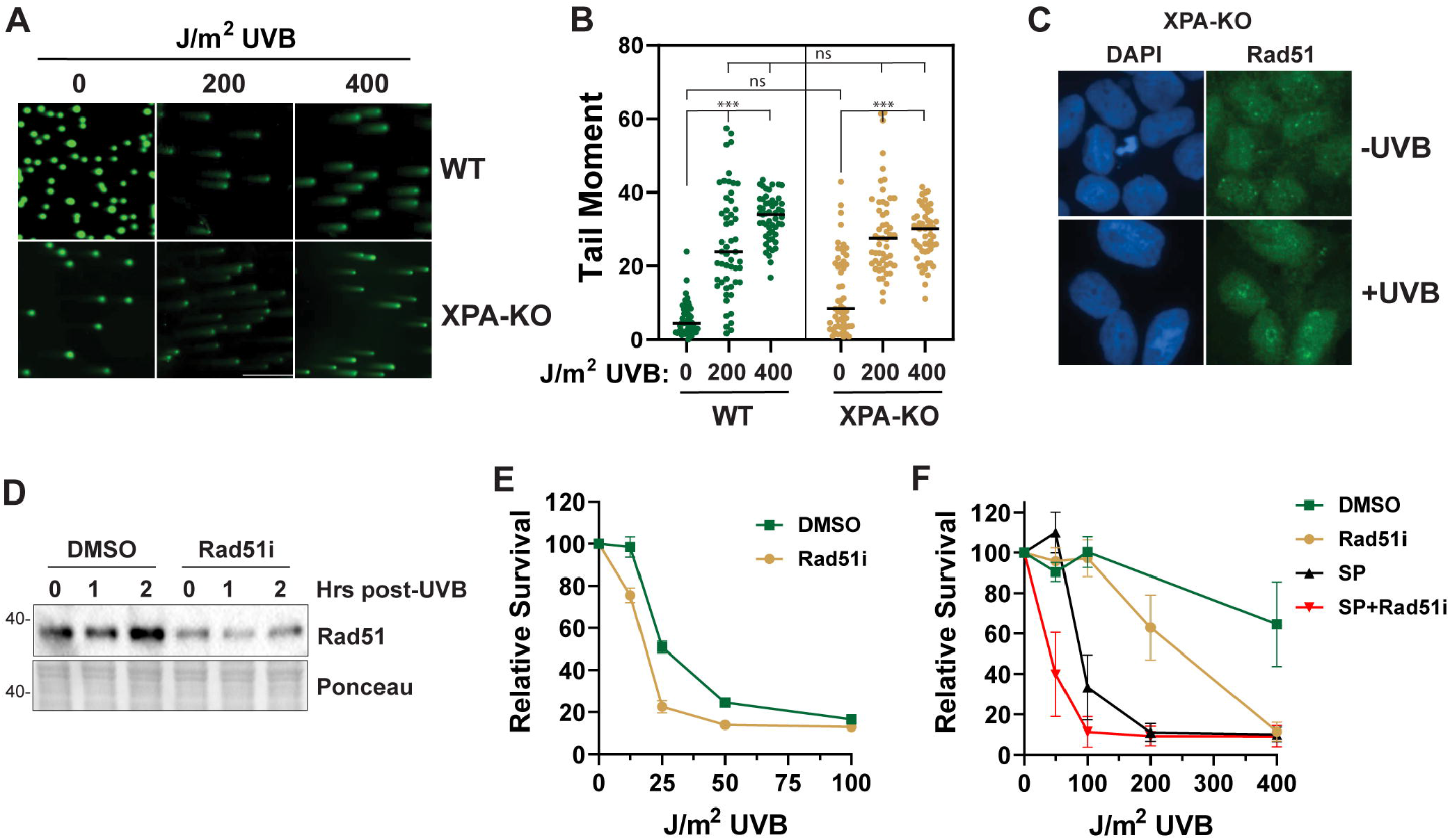
NER-deficient cells exposed to UVB radiation exhibit DNA double-strand break formation and are sensitive to Rad51 inhibition. **(A)** Representative image showing Comet tails in WT and XPA-KO cells exposed to the indicated fluence of UVB radiation. **(B)** Quantitative analysis of Comet tail moment from 50 individual cells analyzed as in (A) with by OpenComet software. An ordinary one-way ANOVA was used to compare treatment groups (***, p<0.001; n.s., not significant). **(C)** Immunofluorescence microscopy was used to examine Rad51 nuclear staining in quiescent XPA-KO HaCaT cells in the absence and presence of UVB exposure. **(D)** XPA-KO cells were treated as indicated and then fractionated to enrich for chromatin- associated proteinss. **(E)** MTT assays were performed 3 days after exposure of quiescent XPA-KO cells treated with DMSO or Rad51i to the indicated fluences of UVB radiation. **(F)** Quiescent HaCaT cells were treated with vehicle or the NER inhibitor spironolactone (SP) in the absence or presence of Rad51i before exposure to the indicated fluences of UVB radiation and measurement of cell survival by MTT assay.

Consistent with the results of neutral Comet assay, we observed that Rad51 formed larger clustered foci by immunofluorescence microscopy (**Fig. 4C**) and accumulated on chromatin (**Fig. 4D**) after UVB exposure. Moreover, treatment of UVB-irradiated quiescent XPA-KO cells with the Rad51 inhibitor resulted in reduced cell viability (**Fig. 4E**). As an additional, independent mechanism of inhibiting NER, quiescent HaCaT cells were treated with spironolactone (SP) for 2 hr to induce the proteolytic degradation of XPB (20, 21), which like XPA is essential for NER. As shown in **Fig. 4F**, Rad51 inhibition sensitized SP-treated quiescent cells to UVB radiation. Thus, we conclude that Rad51 function in promoting cell survival after UVB exposure is independent of NER.

### DNA damage response kinase signaling in UVB-irradiated quiescent cells exhibits both NER- and caspase-dependent and -independent protein phosphorylation events

The ATM protein kinase, which is activated in response to DSBs, has also been reported to be activated in UV-irradiated cells in an NER-dependent manner (Wakasugi et al., 2014). We therefore examined the phosphorylation of ATM and other DNA damage response substrates in both WT and XPA-KO quiescent cells exposed to UVB radiation. As shown in **Fig. 5A** and **Fig. S4**, we observed phosphorylation of all 3 major DNA damage response (DDR) protein kinases (ATR, ATM, and DNA-PKcs) after UVB exposure in both WT and XPA-KO cells. Interestingly, different patterns of phosphorylation were observed with other DDR protein substrates. For example, whereas phosphorylation of the transcriptional regulator KAP1 (Ziv et al., 2006) and histone variant protein H2AX were largely abrogated in XPA-KO cells, only modest effects were observed in the phosphorylation of ATM/ATR target protein p53 (**Fig. S4**), and little effect was found on the phosphorylation of the ATM substrate CHK2. Finally, increased phosphorylation of DNA-PKcs and RPA2 (Ser4/8) were observed 4 hr after UVB exposure in XPA-KO cells in comparison to the WT cells.

**Fig. 5.**
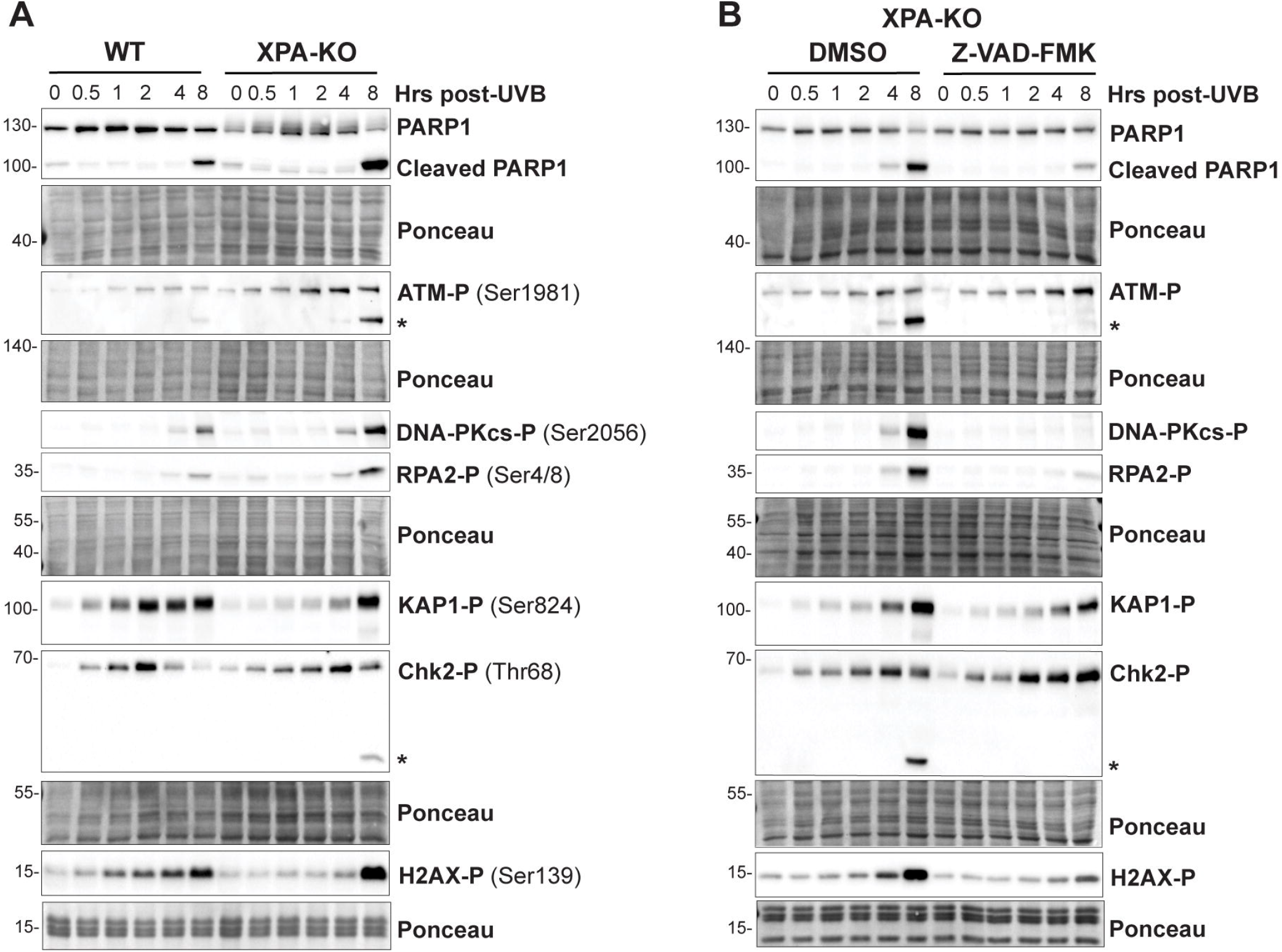
DNA damage response kinase signaling in UVB-irradiated quiescent cells exhibits both NER- and caspase-dependent and -independent protein phosphorylation events. **(A)** Immunoblot analysis of protein phosphorylation in quiescent WT and XPA-KO HaCaT cells after exposure to 200 J/m^2^ UVB. **(B)** Quiescent XPA-KO cells were treated with DMSO or with 20 μM of the pan-caspase inhibitor Z-VAD-FMK before exposure to 200 J/m^2^ UVB. Cells were harvested at the indicated time points and analyzed by immunoblotting. The asterisks indicate likely fragments of ATM and CHK2 generated by caspase signaling.

Because XPA-KO cells are much more sensitive to UVB radiation than WT cells (**Fig. 3C**), we next considered that DSB formation and ATM-dependent protein kinase signaling may be associated with increased or earlier apoptotic signaling following UVB exposure. However, though we observed increased cleavage of the caspase target protein PARP1 in XPA-KO cells relative to WT cells (**Fig. 5A**), this cleavage was only evident 8 hr after UVB exposure in both cell lines and not at the time point (2 hr) at which DSBs were detected (**Fig. 4A**). We noted that the late, 8 hr time point also corresponded to a sudden increase in H2AX phosphorylation in XPA-KO cells (**Fig. 5A**) and with ATM- and CHK2- immunoreactive bands of smaller molecular weight consistent with possible cleavage of these proteins by caspases.

Earlier studies have reported that H2AX can become phosphorylated during apoptosis (Mukherjee et al., 2006; Solier and Pommier, 2009) and that ATM is a caspase target that is inactivated during apoptosis (Smith et al., 1999). To determine whether any of the protein phosphorylation events were associated with apoptotic signaling, we therefore treated XPA-KO cells with the pan-caspase inhibitor Z-VAD-FMK before UVB exposure and then re-examined protein cleavage and phosphorylation events. As shown in **Fig. 5B**, Z-VAD-FMK greatly reduced the level of PARP cleavage and eliminated the apparent cleavage bands for phospho-ATM and phospho-CHK2. Moreover, the elevated phosphorylation of DNA-PKcs, RPA2, and H2AX seen at the 8 hr time point following UVB exposure were all eliminated or greatly reduced by caspase inhibition. We conclude that there are multiple pathways leading to DDR protein phosphorylation in UVB-irradiated quiescent cells and that depending on the specific substrate protein, the phosphorylation may be either dependent or independent of NER and apoptotic signaling. Finally, as previously described (Cleaver, 2011; Marti et al., 2006), these results highlight a limitation of using phospho-H2AX (and DNA-PKcs or RPA2 Ser4/8) as biochemical markers of DSB formation. Nonetheless, we also conclude from these studies that DSB formation and ATM activation in UVB-irradiated cells occurs even in the absence of NER.

### Rad51 promotes survival in response to diverse genotoxic stressors in quiescent cells

Because our data thus far indicated that NER was not required for DSB formation or Rad51 function in quiescent cells, we next explored whether quiescent cells may require Rad51 function in response to exposure to other classes of genotoxins. We first used the RNA splicing inhibitor Pladienolide B, which is known to induce RNA-DNA hybrids (R-loops) in genomic DNA that are potentially prone to breakage and DSB formation (Wan et al., 2015). We observed that quiescent cells were sensitized to Pladienolide when Rad51 function was inhibited (**Fig. 6A**). Similarly, the Rad51 inhibitor sensitized quiescent cells to an inhibitor of RNA polymerase III (Wu et al., 2003), which is required for transcription of 5S rRNA, tRNA, and other small RNAs (**Fig. 6B**). Finally, Rad51 inhibition also sensitized both quiescent HaCaT and U2OS cells to the topoisomerase inhibitor camptothecin (CPT) (**Fig. 6C, D**), which can induce transcription-dependent DSBs in non-replicating cells (Sordet et al., 2010). Indeed, we observed robust phosphorylation of several DDR protein substrates in quiescent cells exposed to CPT (**Fig. 6E**). Analysis of caspase signaling in camptothecin-treated quiescent cells confirmed that Rad51 inhibition potentiates apoptotic signaling (**Fig. 6F**). Thus, we conclude that Rad51 function contributes to the viability of quiescent cells following exposure to many different types of genotoxic agents, including those induce transcription stress.

**Fig. 6.**
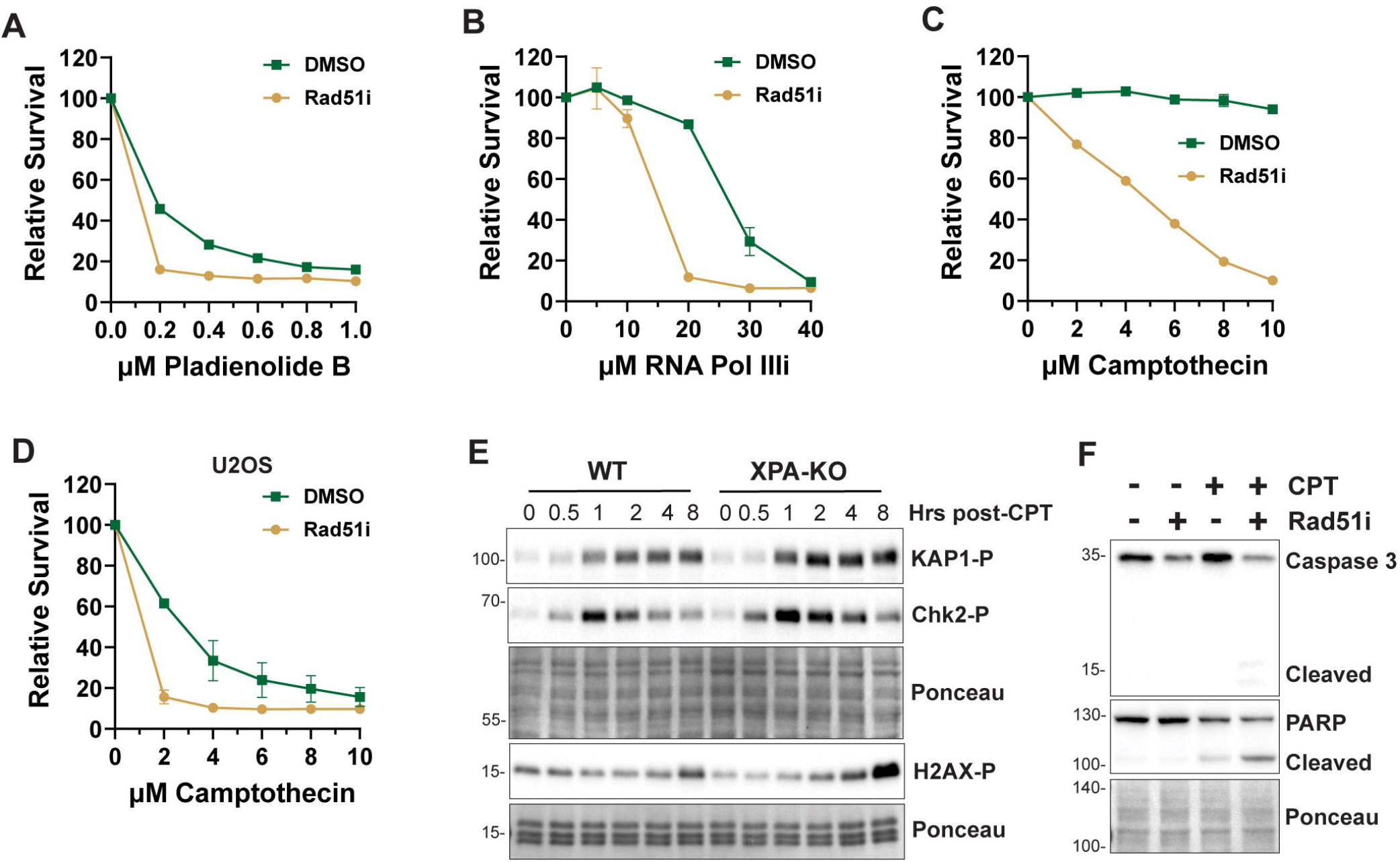
Rad51 inhibition sensitizes quiescent cells to diverse genotoxic agents. **(A)** Quiescent HaCaT cells were treated with the Rad51i and the RNA splicing inhibitor Pladeinolide B, **(B)** the RNA Polymerase III inhibitor ML-60218, or **(C)** the topoisomerase inhibitor camptothecin. MTT assays were performed 3 days later. Results show average and SEM from 2-3 independent experiments. **(D)** Quiescent U2OS cells were treated as in (C). **(E)** Immunoblot analysis of DNA damage kinase signaling in quiescent WT and XPA-KO HaCaT cells treated with CPT for the indicated periods of time. **(F)** Quiescent HaCaT cells were treated with camptothecin (CPT) in the absence or presence of Rad51i for 16 hr before analysis of apoptotic signaling by immunoblotting.

## Discussion

Studies of DNA damage and subsequent cellular responses have traditionally involved the use of model systems in which DNA synthesis and cell cycle progression are major components. Less is generally known about the DNA damage response in non-replicating cells, which comprise the bulk of cells in the human body. Understanding the DNA damage response in this context may provide new insights into how mutagenesis and cell death contribute to cancer, aging, and possibly other disease states.

Here we re-visited a previous report that the generation of DSBs in UVB-irradiated quiescent cells is dependent on NER (Wakasugi et al., 2014) by using CRISPR/Cas9 genome editing to knockout XPA in HaCaT keratinocytes, which have been a useful model system for studying DNA damage responses in quiescent cells (Hutcherson and Kemp, 2019; Kemp, 2017; Kemp and Sancar, 2016; Khan et al., 2022; Shaj et al., 2020). In contrast to the previous study (Wakasugi et al., 2014), we find that DSBs are readily detected in UVB-irradiated cells (**Fig. 4A, B**). We note that there were several differences in the approaches and cell types used by Wakasugi et al and by us here. Whereas Wakasugi et al used a testicular teratoma cell line as their normal, NER-proficient control cell line and fibroblasts from a female patient with a mutant XPA gene, we used CRISPR/Cas9 editing to disrupt the XPA gene in cells with an identical genetic background (HaCaT keratinocytes) (Boukamp et al., 1988). There are several other additional technical differences between the former and current studies, including in the time point post-UV exposure that was analyzed (1 hr vs. 2 hr), the UV light source and dose (20 J/m^2^ UVC vs. 200 J/m^2^ UVB), and the length of the electrophoresis (20 min vs. 30 min). Nonetheless, our observation that inhibition of the DNA recombinase Rad51 sensitizes quiescent XPA-KO and SP-treated cells to UVB (**Fig. 4E, F**) and that robust ATM-CHK2 signaling is found in XPA-KO cells (**Fig. 5**) indicates that DSBs are indeed being generated in the absence of NER. Our further observation that Rad51 sensitizes quiescent cells to several other compounds that directly or potentially impact transcription (**Fig. 6**) suggests that the DSBs that are generated in UVB-irradiated cells may be caused by some form of transcription stress. Nonetheless, future work will be necessary to determine the mechanisms by which DSBs arise in quiescent cells. Finally, our studies making use of a pan-caspase inhibitor (**Fig. 5C**) highlight the potential limitations of using the phosphorylation of H2AX, DNA-PKcs, and RPA2 (Ser4/8) as biochemical markers of DSB formation at time points that correspond to an induction of apoptotic signaling.

Though Rad51 function in DSB repair is primarily thought to occur during S and G2 phases of the cell cycle because of the presence of homologous DNA templates, several recent reports have found that Rad51 can be recruited to DSBs to promote HR under specific experimental conditions in G0/G1. For example, the disruption of proteins and other factors that regulate the expression or stability of HR proteins have both been shown to promote end resection and Rad51 foci formation in G0/G1 (Chen et al., 2021; Orthwein et al., 2015; Xie et al., 2020). Furthermore, studies that have generated DSBs at centromeric regions (Yilmaz et al., 2021) and recruited Cas9 to telomeric DNA to induce ssDNA (Crefcoeur et al., 2017) have shown that Rad51 can be recruited to these specific chromosomal regions characterized by repetitive DNA elements. Finally, the site-specific induction of oxidative stress at a define genomic locus was shown to lead to the recruitment of Rad51 and other HR factors in a transcription- and CSB-dependent manner (Wei et al., 2015).

Thus, our findings here add to this recent data to show that Rad51 plays an important function in the ability of non-replicating quiescent cells to survive exposure to diverse genotoxic insults that include both environmental carcinogens (UVB radiation), cancer chemotherapeutics (cisplatin, camptothecin), and agents that induce transcription stress (Pladienolide B and RNA polymerase III inhibition). Because of the previous reports of Rad51 recruitment to DSBs at centromeres (Yilmaz et al., 2021) and telomeres (Crefcoeur et al., 2017), we expect that Rad51’s pro-survival function in response to DNA damaging agents in quiescent cells may similarly be restricted to specific chromosomal regions and possibly to regions of the genome composed of repetitive DNA elements, which may provide homologous DNA templates for Rad51- mediated HR. Thus, it will be important in future studies to determine whether the location of UV photoproducts and other forms of damage correlate with sites of DSB formation and Rad51 recruitment throughout the genome.

## Materials and methods

### Cell culture

HaCaT and U2OS cells were cultured in DMEM containing 10% FBS and penicillin/streptomycin in a 5% CO_2_ humidified incubator at 37°C. Cells were grown to confluence and then maintained in DMEM containing 0.5% FBS and penicillin/streptomycin for 2-3 days to reach a non-replicating quiescent state. HaCaT cells were transfected with plasmids expressing gRNA targeting XPA and a plasmid with homology direct repair template (Santa Cruz sc-401483) and then single cell clones were selected with puromycin. XPA-KO HaCaT cells stably expressing Flag-tagged XPA were generated by transfecting cells with pcDNA4-Flag-XPA (Addgene #22895) and then selecting single clones with geneticin. Unless otherwise indicated, cells were pre-treated with vehicle (0.1-0.2% DMSO) or the indicated inhibitor for 30 min hr before exposure to UVB radiation (two UVP XX-15M bulbs from Analytik Jena). To inhibit enzymes involved in various DSB repair pathways, cells were treated with the ATM inhibitor (ATMi) KU55933 (10 μM), ATRi VE-821 (10 μM), and DNA-PKi NU7441 (5 μM) (all from Selleckchem) and the PARPi AZD2461 (200 nM), Polθi novobiocin (10 μM), Rad52i DI-03 (10 μM), BLMi ML216 (5 μM), and Rad51i B02 (10 μM) (all from MilliporeSigma). Pladienolide B (1 μM) was from Santa Cruz Biotechnology. With the exception of the Polθi, which was added immediately after UVB exposure due its ability to directly absorb UVB wavelengths, all other compounds were added to the culture medium 30 min before UVB exposure. Acute cell survival was determined using methylthiazolyldiphenyl-tetrazolium bromide (MTT; Sigma), which was added to the cell culture medium at a final concentration of 0.25 mg/ml and incubated for 30 min before solubilization with DMSO and measurement of absorbance at 570 nm on a Synergy H1 spectrophotometer (Bio-Tek). Absorbance values for the UV-irradiated samples were normalized to the non-irradiated samples. Clonogenic cell survival assays involved treatment with the Rad51 inhibitor and UVB, replacement of the medium with fresh, drug-free medium 24 hr later, and then plating 200-500 cells into cell culture plates in normal growth medium 4 days later. Cells were allowed to grow for 10-14 days before staining with crystal violet. Rad51 was knocked down using a pool of Rad51 siRNAs (Santa Cruz sc-36361) and siRNA transfection reagent (Santa Cruz Biotechnology).

### Immunoblotting

Cells were lysed in either RIPA buffer and or a Triton X-100 lysis buffer (20 mM Tris-HCl, pH 7.5, 150 mM NaCl, 1 mM EDTA, 1 mM EGTA, and 1% Triton X-100) for 15-20 min ice and then centrifuged for 20 min at maximum speed in a microcentrifuge at 4°C. Cell lysates were separated by SDS-PAGE, transferred to a nitrocellulose membrane, and then stained with Ponceau S to ensure equal protein loading. Following washing with TBST (Tris-buffered saline containing 0.1% Tween-20) and blocking in 5% milk in TBST, blots were probed with 1:2000 or 1:5000 dilutions of antibodies against XPA (Abcam ab65963), PARP (Cell Signaling #9542), Rad51 (Abcam ab133534), or phosphorylated KAP1 (Ser824; Bethyl A300-767A), p53 (Ser15; Cell Signaling #9248), or H2AX (Ser139; Cell Signaling #9718). After washing, the blots were probed with HRP-coupled anti-mouse or anti-rabbit IgG (ThermoFisher) secondary antibodies for 1-2 hr at room temperature. Chemiluminescence was visualized with either Clarity Western ECL substrate (Bio-Rad) or SuperSignal West Femto substrate (Thermo Scientific) using a Molecular Imager Chemi-Doc XRS+ imaging system (Bio-Rad). Signals in the linear range of detection were quantified by densitometry using Image Lab (Bio-Rad) and normalized to the Ponceau S stained membrane.

### DNA repair and replication assays

For DNA immunoblotting, genomic DNA was purified from treated cells with a Sigma GenElute kit and quantified on a Nanodrop One spectrophotometer. Equal amounts of DNA were boiled for 10 min before cooling on ice and addition of an equal volume of 2 M ammonium acetate. DNA was immobilized on a nitrocellulose membrane and then probed with antibodies against (6-4)PPs (Cosmo Bio NM-DND-002) or ssDNA (Millipore MAB3034). Chemiluminescence was detected as for protein immunoblotting. The excised DNA oligonucleotide products of NER were detected as previously described (Choi et al., 2014; Kim et al., 2022; Song et al., 2017) except that poly-HRP streptavidin was used. To monitor both bulk chromosomal and DNA repair synthesis, proliferating and quiescent WT and XPA-KO cells were exposed or not to 200 J/m^2^ UVB radiation and incubated for 2 hr with 10 μM EdU. Cells were then washed three times with PBS, trypsinized, resuspended in cell culture medium, and pelleted. Cells were then fixed in 4% paraformaldehyde for 15 min, washed once with 1 mL of cold PBS, pelleted, and stored at 4C until staining. Prior to staining, cells were incubated in cold PBS containing 1X saponin and 1% BSA in cold PBS for 15 minutes and subjected to click reactions with Alexa Fluor 647-azide in PBS for 30 min. Cells were analyzed on a Accuri C6 flow cytometer. DNA synthesis associated with S phase replication could be clearly differentiated from cells with lower incorporation of EdU associated with UVB exposure in the WT but not XPA-KO cells.

### DSB detection

DSB formation was detected by neutral comet assay according to the manufacturer’s protocol (Comet assay kit; Trevigen). Briefly, Cells were exposed to 200 J/m^2^ UVB, incubated for 2 hr, and then trypsinized and resuspended in PBS. 1 X 10^5^ Cells were then mixed with molten agarose and transferred onto glass slides followed by incubation in prechilled lysis solution for 1hr to overnight for added sensitivity. The slides were then equilibrated in 1X Neutral Electrophoresis Buffer (NEB) buffer (1 M Tris base, 3 M sodium acetate anhydrous pH 9) for 30 min and electrophoreses at 21 V for 30 min in 1X NEB. Slides were subsequently stained with SYBR Green I and imaged on a Bio-Tek Cytation 5 imager. Comet tail moment was measured in at least 50 cells with OpenComet software (Gyori et al., 2014).

### Immunofluorescence microscopy

Quiescent cells were incubated with 0.25% trypsin-EDTA to dislodge the cells and then treated with defined trypsin inhibitor (Gibco DTI R-007-100). The cells were then seeded onto 12-mm poly-L-lysine coated glass coverslips (Corning 354085, sterilized by UV for 10 min) in a 12-well plate in low serum containing medium. One day later after seeding, cells were irradiated with 200 J/m^2^ UVB and incubated for 2 hr. Cells were then fixed with 4% formaldehyde for 10 min. After several washes with ice-cold PBS, the cells were permeabilized in PBS containing 0.1% Triton X-100 for 10 min at room temperature and followed by a wash with ice cold PBS containing 0.1% Tween-20. Cells were blocked with 5% BSA in PBS for 30 min at room temperature and incubated with anti-Rad51(abcam ab133534) primary antibody at a 1:500 dilution in 1% BSA for 90 min at room temperature or 24 hr at 4°C and was followed by incubation with Alexa Fluor 488 goat anti-rabbit secondary antibody (Invitrogen A11008; 4 μg/ml) in 0.2% BSA for 60 min at room temperature. Cells were then washed twice with 1X PBS. Coverslips were carefully transferred onto glass slides and counterstained with Prolong^®^ Gold antifade with DAPI (Cell Signaling Technologies 8961S). Images were captured at 60 X magnification with a BioTek Cytation 5 Microscope.

## Acknowledgments

We thank the WSU Proteome Analysis Laboratory and Center for Genomics Research for the use of equipment to carry out this work.

## Data Availability

All data are contained within the manuscript.

## CRediT statement

**Saman Khan:** Conceptualization, Methodology, Data Curation, Validation, Writing – Review & Editing; **Alex Carpenter:** Date Curation; **Michael Kemp:** Conceptualization, Supervision, Funding acquisition, Project administration, Writing – Original Draft.

